# Programmed Double-stranded RNA Formation Enables Meiotic Stage Transitions

**DOI:** 10.64898/2026.01.20.700650

**Authors:** Hao Wu, Xinan Liu, Yihan Xiong, Yun Bai, Kevin M. Weeks

**Author notes:** Correspondence (H.W.); (Y.B.); (K.M.W.). These authors contributed equally (Hao Wu, Xinan Liu). Deceased (Yun Bai).

## Abstract

Cell fate transitions rely on extensive transcriptional activation mediated by transcription factors, yet the equally important processes that ensure the selective removal of pre-existing mRNAs remain elusive. We uncover widespread formation of double-stranded RNA (dsRNA) during early-to-middle meiosis using quantitative RNA structure analysis, and validate hundreds of natural antisense transcripts (NATs) through long-read sequencing. These NATs are induced by meiosis-specific transcription factors, including Ndt80p, and pair with sense mRNAs to drive cytoplasmic aggregation of the resulting dsRNAs. These dsRNA aggregates are subsequently transported to the vacuole for clearance, likely via autophagy. This pathway selectively eliminates mRNAs before metaphase I, including *NDJ1*, which encodes a meiotic cohesion protein that would otherwise block chromosome segregation. Our study reveals a physiological role for large-scale dsRNA formation, and highlights a simple but powerful model: the same transcription factors that activate mRNAs required for a specific developmental stage also program the synthesis of NATs that promote removal of transcripts from the preceding stage, thereby driving bidirectional transcriptome reprogramming.

## INTRODUCTION

The transcriptome represents a balance between mRNA transcription and degradation^1^. During major cell transitions, the composition of the transcriptome undergoes dramatic changes over short time periods. The production phase is orchestrated by a transcriptional cascade^2^, in which transcription factors and regulatory networks sequentially activate distinct sets of genes with high specificity. In contrast, major RNA degradation pathways for RNA polymerase II transcripts in eukaryotes, including nuclease-dependent decay and translation-associated decay^3^, generally operate with little sequence specificity. This discrepancy highlights a fundamental gap in our understanding of how cells precisely and temporally eliminate pre-existing mRNAs, which differ widely in stability and abundance, during cell fate transitions.

Antisense transcription occurs across diverse biological processes^4,5^. The resulting products, known as natural antisense transcripts (NATs), undergo typical RNA processing steps including 5′-capping and 3′-polyadenylation, and can be exported to the cytoplasm^6^. Traditionally, NATs have been viewed as regulators of gene expression through transcriptional interference^7^ or RNA interference (RNAi)^8^. A productive role for cytoplasmic NATs in forming long double-stranded RNAs (dsRNAs) with sense transcripts has generally been overlooked, as dsRNAs have primarily been viewed as hallmarks of viral infection that activate the interferon signaling pathway^9^. More recent studies, however, have uncovered physiological functions for dsRNA formation, including promoting nuclear export of sense mRNAs^10^ or repressing their translation^11^. Despite the prevalence of antisense transcription, the conditions under which dsRNAs form and their broader biological functions are not well understood.

Meiosis is a tightly regulated developmental program characterized by multiple large-scale transitions between successive stages. Meiotic prophase I, in particular, is governed by checkpoint mechanisms, in which key proteins produced in earlier stages prevent premature entry into subsequent stages, therefore ensuring accurate chromosome segregation^12,13^. Progression through these transitions requires active suppression of checkpoint proteins through diverse regulatory strategies, including targeted protein degradation, post-translational modifications, and changes in subcellular localization^12,13^. Equally critical is the clearance of mRNAs whose persistent translation would interfere with accurate chromosome segregation, though the underlying mechanisms are poorly understood. Antisense transcription occurs during meiosis across diverse species, from fungi to mammals^14,15^, and dsRNA formation has been detected in budding yeast *S. cerevisiae*^16^ and mouse testes^17^. However, the functional significance of dsRNA formation during meiosis remains unclear. Notably, *S. cerevisiae* lacks the canonical RNAi machinery found in many eukaryotes^18^, which processes dsRNA into small interfering RNAs. This absence makes yeast an ideal model for dissecting alternative dsRNA-based regulatory mechanisms.

In this study, we identified extensive changes in mRNA structure during early-to-middle meiosis through time-resolved dimethyl sulfate modification analyzed by mutational profiling (DMS-MaP). Using a dsRNA-specific antibody, we detected large-scale dsRNA formation during this period and observed its assembly into cytoplasmic aggregates. We further identified hundreds of meiosis-specific NATs from time-resolved, strand-specific RNA-seq. The double-stranded structures formed between sense transcripts and NATs promote their aggregation, as artificially expressed *TIP1* antisense RNA (*TIP1-AS*) co-localized with the dsRNA aggregates. Motif enrichment analysis, together with nanopore sequencing, identified the meiosis-specific transcription factors Ume6p and Ndt80p as stage-specific activators of antisense transcription via the *URS1* (Upstream Repressing Sequence 1) and *MSE* (Middle Sporulation Element) *cis*-regulatory DNA motifs^19^. Moreover, we performed co-localization imaging with markers for P-bodies, stress granules, autophagosomes, and vacuole membrane to explore the functional role of dsRNA formation. We observed that dsRNA aggregates are transported to the vacuole for degradation, most likely via autophagy, as large dsRNA granules co-localized with the autophagy marker *ATG8*. This mechanism facilitates synchronized removal of stage-conflicting mRNAs, including *NDJ1*, which encodes a cohesion protein that prevents premature chromosome segregation^20^. Specifically, a pair of sequentially activated, convergent promoters at the *NDJ1* locus program the timing of dsRNA formation, shaping its pulsed, stage-specific expression during meiotic prophase. Together, our findings demonstrate that programmed dsRNA formation functions as a large-scale, yet selective, strategy for transcript clearance that promotes precise meiotic stage transitions.

## RESULTS

### Time-resolved Structure Profiles Reveal mRNA Structural Rearrangements During Meiosis

In a previous study, we generated comprehensive mRNA structure profiles across synchronized yeast meiosis at nine stages^21^ – vegetative growth (V), pre-meiotic (P), pachytene (T0), end of prophase I (T1), metaphase I (T2), anaphase I (T3), metaphase II (T4), anaphase II (T5), and meiotic exit (T6) – using dimethyl sulfate mutational profiling (DMS-MaP) (Fig. 1A and Fig. S1A). DMS-MaP quantitatively measures chemical accessibility of DMS to RNA at single-nucleotide resolution, termed DMS reactivity. DMS reactivity changes across stages thus reflect RNA structural dynamics throughout meiosis. Although translation is extensively regulated during yeast meiosis^22^, our earlier work showed that global mRNA structural remodeling occurs largely independent of translation^21^, suggesting that major RNA structural changes are driven by other meiotic processes.

**Figure 1.**
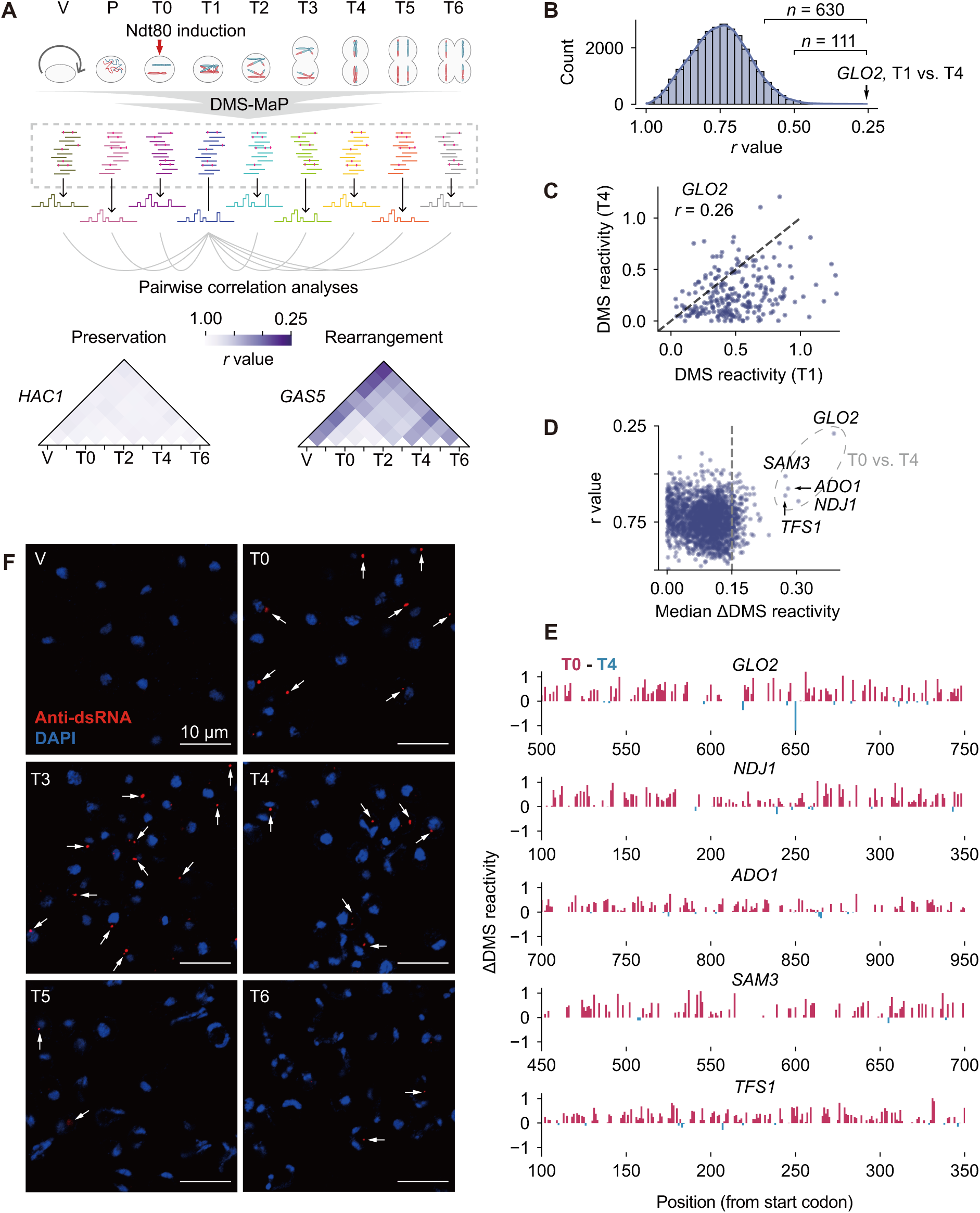
**RNA structure dynamics analyses reveal dsRNA formation during early-to-middle meiosis** (A) Overview of RNA structural dynamics during meiosis. Pairwise Pearson correlation coefficients (*r*) of DMS reactivity profiles were calculated between all meiotic stages. Lower *r* values indicate greater structural remodeling. Representative examples illustrate highly preserved structure (*HAC1*) and pronounced stage-specific rearrangement (*GAS5*). The DMS-MaP dataset was generated in our prior study^21^. (B) Distribution of Pearson correlation coefficients (*r*) from 28,601 pairwise comparisons among 2,058 mRNAs. *n* indicates the number of mRNAs included. Low r-values correspond to large RNA structural differences between states. (C) Comparison of individual per-nucleotide DMS reactivity measurements for *GLO2* between the T0 and T4 stages. (D) Median ΔDMS reactivity of 28,601 pairwise comparisons and their corresponding *r* values. The five genes highlighted by the dashed circle exhibit the largest DMS reactivity changes between T0 and T4 states. (E) DMS reactivity profiles of 250-nt regions within the *GLO2, NDJ1, ADO1, SAM3,* and *TFS1* mRNAs, comparing the T0 and T4 states. High ΔDMS values correspond to reduced reactivity in T4. (F) Immunofluorescence staining of dsRNA during meiosis. Red: anti-dsRNA antibody indicating dsRNA; blue: DAPI staining showing nuclei. As the time intervals between T1, T2, and T3 were very close (T1: 7 h; T2: 7.25 h; T3: 7.5 h), only the T3 sample was analyzed here. White arrows indicate dsRNA signals.

In this study, we used DMS-MaP data to identify mRNAs undergoing the most pronounced structural remodeling and to determine their temporal occurrence. We quantified structural similarity between pairwise stages by Pearson correlation of DMS reactivities (Fig. 1A). For each transcript, the lowest correlation coefficient (*r*) indicates the stage pair with the greatest structural change. Among transcripts with sufficient coverage across all stages, we observed markedly different remodeling patterns. Some transcripts, like *HAC1* and *SOD1*, maintained consistent structures throughout the entire meiotic process (minimum *r* > 0.88; Fig. 1A and Fig. S1B). In contrast, others underwent notable stage-specific alterations. For example, *GAS5* retained stable folding from the pre-meiotic (P) stage to metaphase I (T2; *r* = 0.84–0.93), but showed dramatic structural changes between vegetative growth (V) and meiotic exit (T6; *r* = 0.41; Fig. 1A). Similarly, *CYT1* exhibited its most pronounced structural change between anaphase I (T3) and meiotic exit (T6; Fig. S1B). These results demonstrate that large-scale RNA structural remodeling is temporally regulated and transcript-specific during meiosis.

We monitored 28,601 pairwise comparisons across 2,058 mRNAs with sufficient coverage. This analysis revealed widespread structural remodeling during meiosis, with 630 genes showing a minimum r value below 0.6, and 111 genes below 0.5 (Fig. 1B and Table S1). The most extreme case is *GLO2* mRNA, which has the lowest r value of 0.26 between prophase I (T1) and metaphase II (T4; Fig. 1B), indicating extensive structural rearrangements during this period. Intriguingly, the majority of nucleotides in *GLO2* showed reduced DMS reactivities at metaphase II (T4) relative to prophase I (T1; Fig. 1C). This one-way shift in reactivity differs from typical intramolecular structural remodeling, which typically involves both increases and decreases in reactivity, reflective of local transitions between single-stranded and double-stranded states.

To identify additional transcripts showing similar one-way changes in reactivity, we calculated per-nucleotide ΔDMS reactivity values between pairwise stages, and used the median to quantify the unidirectional remodeling for each transcript. This analysis identified 245 genes with median ΔDMS reactivity exceeding 0.15 (Fig. S1C and Table S1). Among these, *GLO2*, *NDJ1*, *ADO1*, *SAM3*, and *TFS1* showed the greatest changes, with median ΔDMS reactivities ranging from 0.27 to 0.39, accompanied by low r values between stages (Fig. 1D). In all five cases, DMS reactivity decreased markedly during the pachytene (T0) to prophase I (T1) transition (Fig. S1D) and reached maximal reduction at metaphase II (T4; Fig. 1D and Fig. S1D). Inspection of individual reactivity profiles confirmed broad decreases in DMS reactivity, across large segments of these mRNAs, at metaphase II (T4) compared with pachytene (T0; Fig. 1E). Collectively, these analyses highlight pervasive RNA structural remodeling during meiosis and indicate that the most dramatic unidirectional rearrangements occur in a specific subset of transcripts during the early-to-middle meiosis.

### Stage-specific NATs Trigger Double-stranded RNA Formation and Aggregation

We attribute the observed one-way reduction in DMS reactivity to extensive intermolecular base-pairing by eliminating the following conventional explanations. (*i*) Intramolecular structural remodeling typically produces bidirectional DMS reactivity changes across different regions^23^. (*ii*) RNA-binding protein interactions generally reduce DMS reactivity locally at their binding sites^24^. (*iii*) Although ribosome scanning and elongation can globally unfold mRNAs, our previous work showed no correlation between RNA structural changes and translational efficiency ^21^. Thus, we considered the hypothesis that the broad, long-range reactivity decreases reflect widespread formation of intermolecular double-stranded RNA (dsRNA). To test this, we stained cells with the J2 antibody, which specifically recognizes dsRNA helices over 40 bp^25^, across multiple stages from vegetative growth (V) to meiotic exit (T6). Consistent with our hypothesis, dsRNAs were detected exclusively during early-to-middle meiosis from pachytene (T0) to metaphase II (T4; Fig. 1F). Strikingly, these dsRNAs assembled into large cytoplasmic, granular structures at pachytene (T0), which began to dissipate by metaphase II (T4) and were completely absent by meiotic exit (T6; Fig. 1F). These observations support the model that the one-way reduction in DMS reactivity observed during early-to-middle meiosis arises from transient, large-scale dsRNA formation.

Given that antisense transcription is evolutionarily conserved during meiosis from yeast to humans^4,14^, we hypothesized that dsRNAs are formed by base pairing between NATs and their sense mRNAs. However, a stage-specific NAT profile in synchronous meiosis had not been established. Leveraging the strand-specificity of our time-resolved RNA-seq libraries, we mapped reads to both strands for each gene locus and calculated antisense read ratios across meiosis (Table S2). Hierarchical clustering of 405 genes with >20% change in antisense read ratio revealed a dynamic NAT landscape during the transition from vegetative growth to meiosis (Fig. 2A). For example, *IME4*, whose antisense transcription determines cell fate through transcriptional interference^7^, showed a high antisense read ratio (48.0%) at the pre-meiotic stage (P) that decreased sharply by pachytene (T0; Fig. 2A and Fig. S2A). However, the majority (313 genes, 77%) exhibited increased antisense read ratios only after the pre-meiotic (P) stage (Fig. 2A), indicating meiosis-specific activation of antisense transcription. Importantly, the five transcripts showing the most pronounced unidirectional structural remodeling in early-to-middle meiosis, including *GLO2*, *NDJ1*, *ADO1*, *SAM3*, and *TFS1* (Fig. 1D), all displayed meiosis-specific activation of antisense transcription (Fig. 2A). These findings strongly support that meiosis-specific NATs drive dsRNA formation during meiosis.

**Figure 2.**
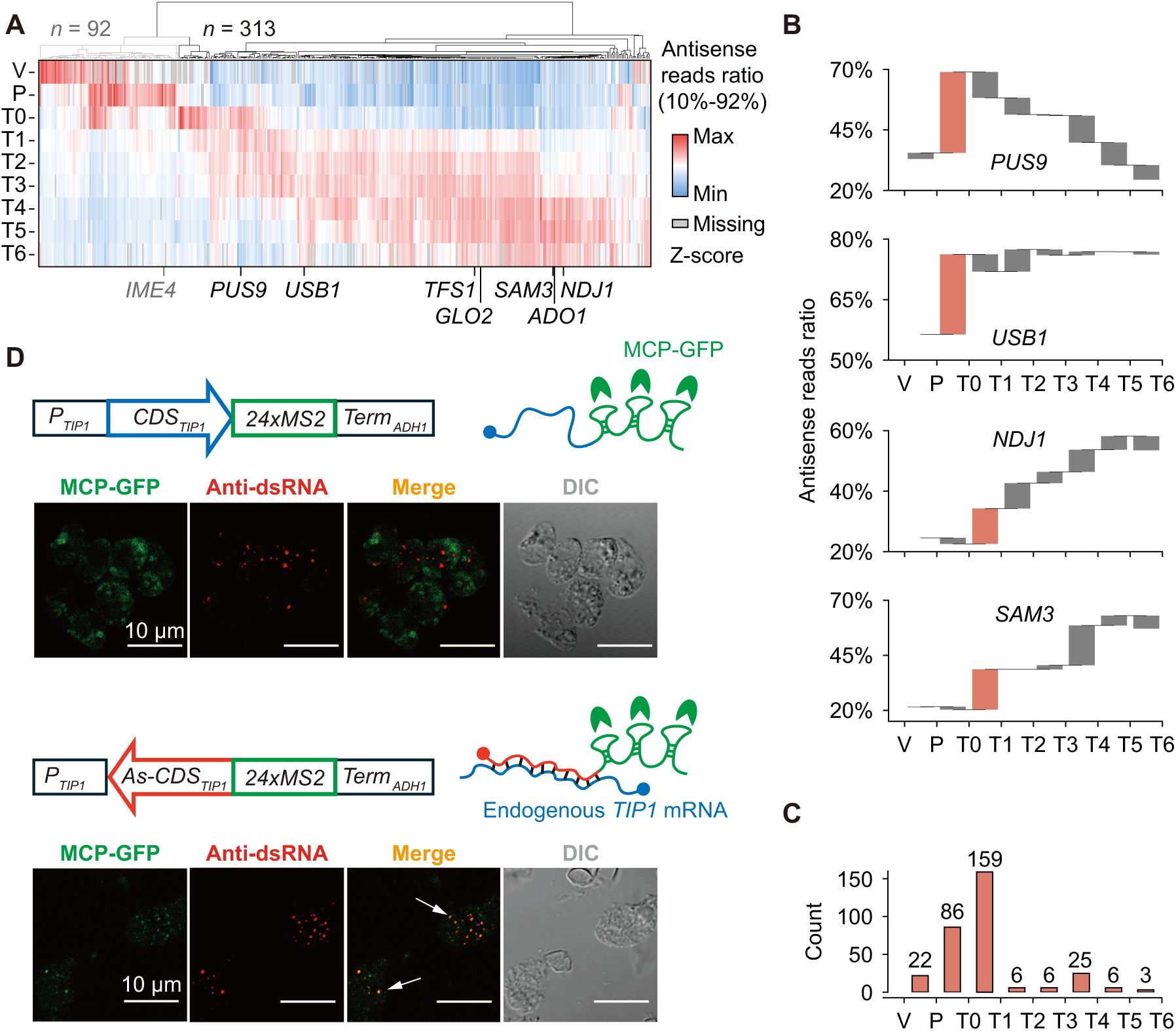
**Meiosis-specific NATs trigger dsRNA aggregation** (A) Hierarchical clustering of 405 genes with >20% change in antisense read ratio. Of these, 313 genes show meiosis-specific increase (from T0 onward) in antisense read ratios. (B) Antisense read ratio dynamics for selected genes (*PUS9*, *USB1*, *NDJ1*, and *SAM3*) throughout meiosis. Intervals of maximal increase are highlighted in orange. (C) Distribution of the 313 genes associated with meiosis-specific NATs according to the intervals of maximum increase in antisense read ratio. (D) Subcellular localization of *TIP1* sense mRNA (top) and artificially expressed *TIP1-AS* (bottom), tracked using the MCP-MS2 system. Green: MCP-GFP indicating sites of MS2-tagged RNA; red: anti-dsRNA antibody indicating positions of dsRNA. White arrows indicate co-localization between dsRNA signals and MCP-GFP.

To further classify the activation timing of meiosis-specific NATs, we defined their activation stage as the interval with the largest increase in antisense read ratio (Table S3). For example, *PUS9* and *USB1*, two genes involved in small RNA processing, showed 20-30% increase in antisense read ratio between the pre-meiotic (P) stage and pachytene (T0; Fig. 2B), while the antisense read ratio for *NDJ1* and *SAM3* began to increase only after pachytene (T0; Fig. 2B). Time-resolved northern blot analysis of NATs from these four genes confirmed that their expression timing was consistent with assigned activation stages (Fig. S2B), supporting the use of antisense read ratio as a proxy for antisense transcription strength. A statistical analysis for all 313 genes with meiosis-specific NATs revealed that 78% are activated within two major intervals: pre-meiotic (P) to pachytene (T0), and pachytene (T0) to prophase I (T1; Fig. 2C). These results indicate that, although NATs are predominantly activated during early-to-middle meiosis, their activation timing remains distinctly stage-specific.

To validate the relationship between NATs and dsRNA aggregates, we introduced an artificial antisense RNA for *TIP1*, a randomly selected gene unrelated to meiosis and lacking endogenous antisense transcription at all stages (Fig. S2C). Using the endogenous *TIP1* promoter, we expressed partial sense *TIP1* mRNA, starting from the 5’ UTR and including the entire coding region, fused to MS2 stem-loop elements at the 3′ end. RNA localization was tracked through MS2 coat protein fused to green fluorescent protein (MCP-GFP). Expression of the *TIP1* sense mRNA alone did not lead to co-localization with dsRNA aggregates at pachytene (T0; Fig. 2D). In contrast, when the *TIP1* sequence was expressed in the reverse complementary orientation to generate *TIP1* antisense RNA (*TIP1-AS*), *TIP1-AS* formed cytoplasmic foci that co-localized with dsRNA aggregates at pachytene (T0; Fig. 2D). These results confirm that dsRNAs undergo selective aggregation and provide direct evidence that NATs are sufficient to trigger dsRNA aggregation.

### Stage-specific Transcription Factors Activate Pervasive Antisense Transcription

We hypothesized that the downstream regions at the 3′ end of these loci should harbor promoters that drive antisense transcription. We then performed de novo motif analysis on the 3′ end sequences of 313 gene loci associated with meiosis-specific NATs. The same regions from all other genes were used as background for motif enrichment analysis, and the enriched motifs were compared to motif databases to identify involved transcription factors (Fig. 3A). Separate analyses for antisense promoters activated in the P-T0 and T0-T1 intervals identified distinct top motifs: 5’-CGGCGGCTAA-3’ and 5’-CGCGGCACAA-3’, respectively (Fig. 3A and Table S4). The 5’-CGGCGGCTAA-3’ motif for the P-T0 interval matches the Upstream Repressing Sequence 1 (URS1) consensus motif (5’-TCGGCGGCTA-3’), while 5’-CGCGGCACAA-3’ for the T0-T1 interval is part of the middle sporulation element (MSE, 5’-GNCRCAAAW-3’; Fig. 3B). The fourth most enriched motif for the T0-T1 interval corresponds exactly to the MSE (Fig. S3A). Importantly, the URS1 and MSE elements are binding sites for Ume6p and Ndt80p, respectively (Fig. 3B), two transcription factors that are essential for meiosis. Ume6p, together with Ime1p, initiates the expression of early meiotic genes required for meiotic entry, whereas Ndt80p, a well-characterized meiosis-specific transcription factor, activates middle meiotic genes to promote the transition from prophase I to chromosome segregation. These findings support the striking finding that meiosis-stage-specific transcription factors drive pervasive antisense transcription through *cis*-regulatory elements located at the 3′ ends of gene loci.

**Figure 3.**
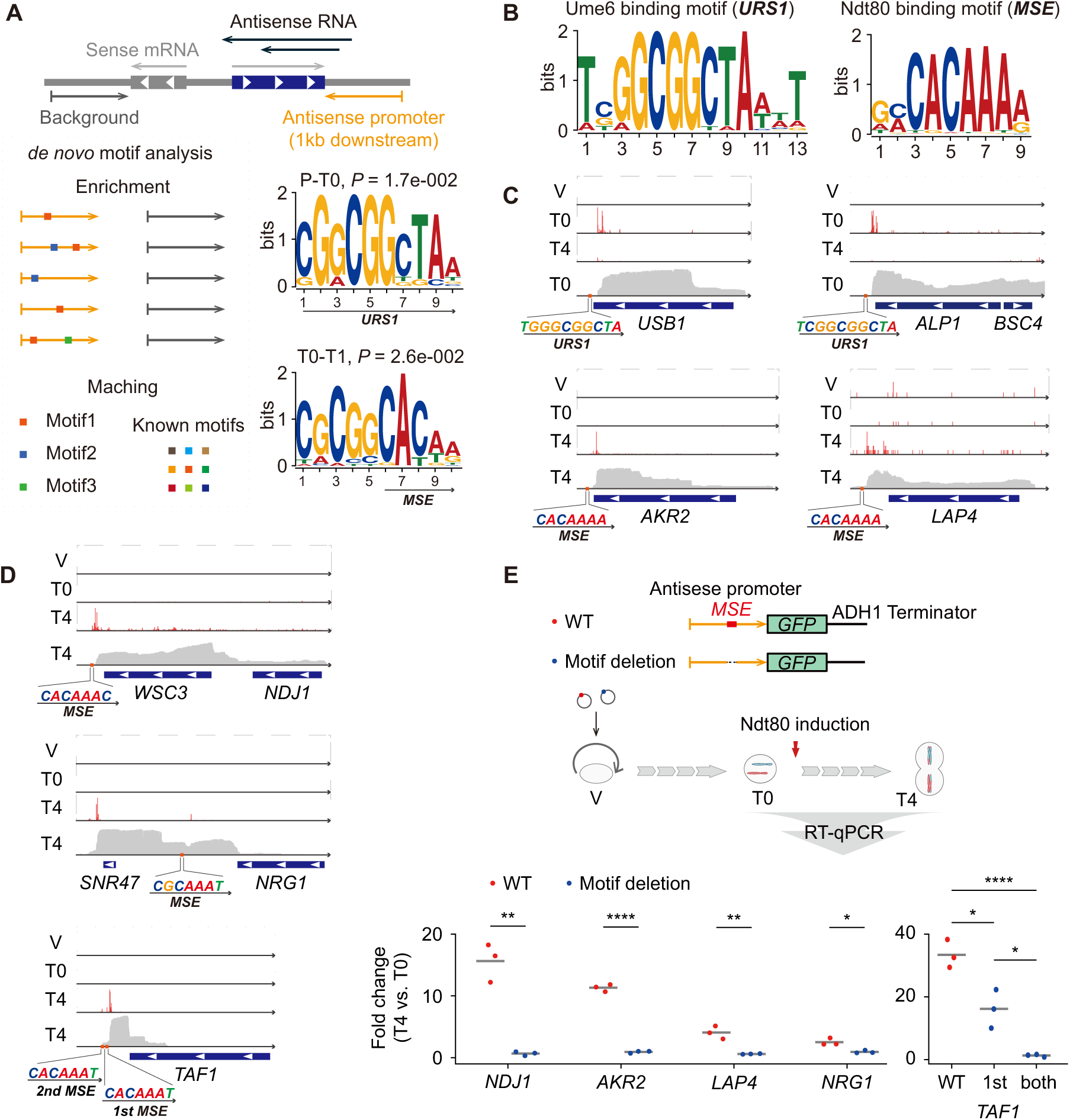
**Stage-specific transcription factors activate NATs via upstream promoter elements** (A) Motif enrichment analysis of antisense promoters. The two top-enriched motifs for antisense promoters with activation stages assigned to the P-T0 and T0-T1 intervals are shown. (B) Consensus Ume6p binding motif (URS1) and Ndt80p binding motif (MSE) obtained from the JASPAR database^45^. (C) Genomic context of URS1 and MSE elements relative to antisense transcription initiation sites for *USB1*, *ALP1*, *AKR2*, and *LAP4*. The transcription initiation sites were defined by 5′-end mapping of long-read sequencing data. (D) Genomic context of MSE elements relative to antisense transcription initiation sites for *NDJ1*, *NRG1*, and *TAF1*. (E) Functional requirement of MSE for antisense promoter activation. Wild-type (red) or core sequence-deleted (5’-*CACAAA*-3’; blue) antisense promoters of the indicated genes (*NDJ1*, *AKR2*, *LAP4*, *NRG1*, and *TAF1*) were inserted upstream of the GFP ORF. Cells were harvested before (T0) and after (T4) Ndt80 induction. Fold changes in GFP mRNA (T4 vs. T0; normalized to *ACT1*) were measured by RT-qPCR. Wild-type promoters yielded 2.7- to 33.4-fold induction. Deletion of a single (for *NDJ1*, *AKR2*, *LAP4* and *NRG1*) or both (for *TAF1*) MSEs (blue points) abolished transcriptional activation to near-background levels (0.6- to 1.3-fold).

To validate the relationship between antisense transcription initiation and *cis*-regulatory elements, we performed full-length cDNA nanopore sequencing at three key stages: vegetative growth (V), pachytene (T0), and metaphase II (T4), achieving an average sequencing depth of 10 million reads per time point. By using RNA samples with high integrity and quantifying the 5′ ends of each read, we accurately mapped transcription initiation sites (Fig. S3B). For example, *HOP1*, a core component of the synaptonemal complex, showed high counts downstream of a URS1 element at the 5’ end of its locus at pachytene (T0), indicating active transcription at early meiosis (Fig. S3C). *SPS4*, a sporulation-specific transcript induced by Ndt80p, exhibited transcription initiation downstream of a MSE at the 5’ end of its locus only at middle meiosis (T4; Fig. S3C). As expected, antisense transcription initiation frequently occurs within 100 bp downstream of the URS1 and MSE elements located at the 3′ ends of gene loci (Fig. 3C). For example, *USB1* and *ALP1* initiate antisense transcription downstream of the URS1 motif during pachytene (T0), while *AKR2* and *LAP4* initiate antisense transcription downstream of the MSE motif only at metaphase II (T4; Fig. 3C). We also identified more complex initiation patterns mediated by MSEs. For example, antisense transcription initiated by an MSE at the *NDJ1* 3′ end spans across the neighboring *WSC3* locus, generating a NAT that covers both genes, indicative of long-range bicistronic regulation (Fig. 3D). At the *NRG1* locus, MSE-driven initiation occurs within the body of another highly transcribed gene (Fig. 3D). In the case of *TAF1*, a subunit of the TFIID transcription factor complex, two tandem MSE motifs may act cooperatively to enhance antisense transcription strength (Fig. 3D). Together, these findings reveal intricate structural relationships between *cis*-regulatory elements and antisense transcription, underscoring the complexity of regulation based on antisense transcription.

To directly validate the stage-specific activation of downstream NATs by *cis*-regulatory elements, we cloned five native antisense promoters (*NDJ1*, *AKR2*, *LAP4*, *NRG1* and *TAF1*), containing the MSEs, into expression vectors and generated variants in which the core MSE sequence (5’-CACAAA-3’) was deleted (Fig. 3E). We then compared downstream transcript levels before and after Ndt80p induction (T4 vs. T0) by reverse transcription quantitative PCR (RT-qPCR; Table S5). All wild-type antisense promoters robustly activated downstream transcripts by 2.7- to 33.4-fold in response to Ndt80p induction (T4 vs T0; Fig. 3E). Deletion of a single MSE in antisense promoters of *NDJ1*, *AKR2*, *LAP4*, and *NRG1* reduced transcriptional activation to near-background levels (0.6- to 1.1-fold; Fig. 3E). Notably, *NDJ1*, which encodes a meiotic cohesion protein that prevents chromosome segregation, exhibited particularly strong stage-specific antisense transcription of 16-fold driven by a single MSE. As for *TAF1*, whose antisense promoter contains two tandem MSEs, stage-specific activation reached 33-fold after Ndt80p induction. Deletion of one MSE reduced activation to 16-fold, while deletion of both completely abolished stage-specific induction (1.3-fold; Fig. 3E), indicating that tandem MSEs enhance the strength of antisense transcription. Collectively, these results validate that *cis*-regulatory elements within antisense promoters mediate activation of NATs, accounting for their temporal specificity and supporting a model in which meiotically essential transcription factors simultaneously induce both mRNAs and NATs during early-to-middle meiotic transitions.

### Double-stranded RNAs Are Detained in Vacuoles

To determine how dsRNA formation affects sense transcripts, we analyzed abundance changes in 313 sense mRNAs associated with meiosis-specific NATs. On average, these mRNAs reach peak abundance during the pre-meiotic (P) stage and undergo a pronounced decrease starting at pachytene (T0; Fig. S4A), a pattern consistent with the canonical role of antisense transcription in suppressing gene expression by transcriptional interference. Unexpectedly, hierarchical clustering of sense mRNA abundance revealed two clusters that displayed pulsed expression — a rapid increase followed by an equally rapid decrease — specifically during the pre-meiotic (P) to prophase I (T1) interval (Fig. 4A), coinciding with the window of antisense transcription (Fig. 2C). This pattern is markedly different from genes whose abundance rebounds after prophase I (T1; Fig. 4A) and genes upregulated during meiosis but lacking antisense transcription (Fig. S4B). These findings suggest that dsRNA formation shapes the pulsed expression profiles during the pre-meiotic (P) to prophase I (T1) transition, supporting a post-transcriptional mechanism that rapidly and synchronously clears stage-specific mRNAs.

**Figure 4.**
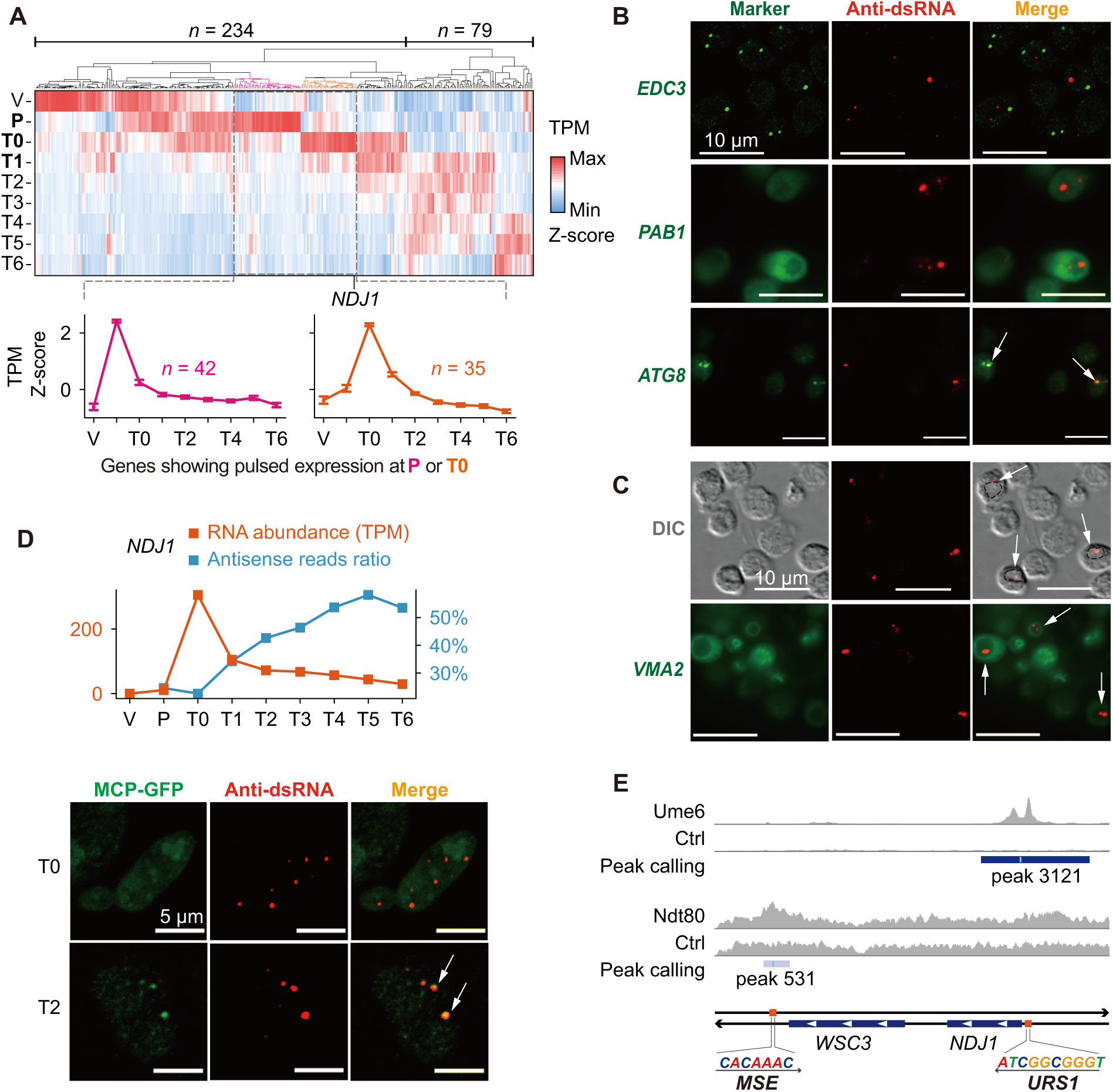
**Vacuolar degradation shapes pulsed gene expression** (A) Top: Hierarchical clustering of sense mRNA abundance for 313 genes with meiosis-specific NATs. Bottom: Average z-scored TPM for two clusters showing pulsed expression at pre-meiotic (P; pink; *n* = 42) and pachytene (T0; orange; *n* = 35) stages. (B) Co-localization of dsRNA aggregates with RNA degradation-related cellular structures. Green: GFP-tagged marker proteins including Edc3-GFP (P-body), Pab1-GFP (stress granule), and GFP-Atg8 (autophagosome); red: anti-dsRNA antibody indicating dsRNA. White arrows indicate co-localization of Atg8 with dsRNA. (C) Top: DIC imaging showing concave cytoplasmic regions and dsRNA aggregates (red) in cells. Dashed circles in the merge channel highlight concave cytoplasmic regions containing dsRNA aggregates. Bottom: Co-localization of dsRNA aggregates (red) with the vacuolar membrane marker Vma2-GFP (green). White arrows indicate dsRNA signals within the vacuole. (D) Top: *NDJ1* sense mRNA abundance (orange line) and antisense read ratio (blue line) across meiosis. Bottom: Co-localization of *NDJ1-MS2* with dsRNA aggregates before (T0) and after (T2) Ndt80p induction. White arrows indicate co-localization between dsRNA signals and MCP-GFP. (E) ChIP-seq occupancy of Ume6p (URS1) and Ndt80p (MSE) at the *NDJ1* locus.

To determine whether dsRNA formation is associated with canonical RNA degradation pathways, we performed co-localization analyses for dsRNA aggregates and GFP-tagged marker proteins for P-bodies (*EDC3*), stress granules (*PAB1*), and autophagosomes (*ATG8*). We found that Edc3-GFP did not co-localize with dsRNA aggregates, indicating that dsRNA aggregates are not degraded via P-bodies (Fig. 4B). Pab1-GFP was diffusely distributed throughout the cell and did not form granules, suggesting the lack of stress granule assembly during meiosis (Fig. 4B). In contrast, large dsRNA aggregates overlapped with GFP-Atg8 (Fig. 4B), implicating autophagy as a potential route for their degradation.

The terminal destination of autophagy, including RNAphagy, is the vacuole^26^, a lysosome-like organelle in yeast. We therefore examined the localization of dsRNA aggregates by differential interference contrast (DIC) microscopy and found that large dsRNA granules occupied punctate sites enclosed within concave cytoplasmic regions typical of vacuolar morphology (Fig. 4C). GFP tagging of the vacuolar membrane protein Vma2 confirmed that large dsRNA aggregates reside inside the vacuole (Fig. 4C). The primary vacuolar RNase responsible for RNA degradation is Rny1^27^. We therefore assessed Rny1 involvement by querying published proteomic data of natural meiosis^28^. We found that Rny1 is highly active throughout meiosis, with protein abundance increasing more than threefold from pachytene (T0) compared to the vegetative (V) stage and continuing to rise thereafter (Fig. S4C), indicating enhanced RNase activity in the vacuole. Collectively, our findings support a model in which dsRNA aggregates are transported via autophagy and ultimately detained in the vacuole for clearance.

### Genetic Circuits that Shape Pulsed Gene Expression

To determine how stage-specific NATs influence their sense transcripts through dsRNA-mediated decay, we focused on two genes, *NDJ1* and *AKR2*, whose antisense promoters showed the strongest MSE-dependent induction. *NDJ1* encodes a meiotic cohesion protein that prevents premature chromosome segregation, while *AKR2* encodes a palmitoyltransferase with an uncharacterized meiotic role. Joint analysis of mRNA abundance and antisense read ratio revealed the pulsed expression of both genes with a sharp decline in mRNA levels concurrent with the initial rise in antisense read ratio (Fig. 4D and Fig. S4D). We tagged these transcripts with MS2 stem-loops and monitored their subcellular localization before and after Ndt80 induction. At pachytene (T0), prior to Ndt80-dependent activation of their NATs, neither transcript formed cytoplasmic foci. In contrast, at metaphase I (T2), both transcripts showed robust co-localization with large dsRNA aggregates (Fig. 4D and Fig. S4E), which we showed to be ultimately transported to the vacuole (Fig. 4C). Together, these results indicate that stage-specific NATs promote the vacuolar clearance of their corresponding sense mRNAs.

Although antisense transcription is widespread across biological systems, dsRNA formation is not commonly observed, likely because it requires simultaneous transcription of both sense and antisense strands, which is energetically inefficient. We propose that dsRNA formation is most likely to occur during phases of cellular transition, when the relative strengths of sense and antisense transcription shift. As *NDJ1* encodes a cohesion protein that controls chromosome segregation, we examined how its pulsed, stage-specific expression is encoded. Interestingly, motif scanning revealed a URS1 element in its sense promoter (Fig. 4E). Since Ume6p (URS1 regulator) functions in early meiosis and Ndt80p (MSE regulator) functions in mid-meiosis, the *NDJ1* locus harbors a pair of temporally ordered, convergent promoters. Analyses of chromatin immunoprecipitation followed by sequencing (ChIP-seq) datasets^29,30^ confirmed binding of the transcription factor Ume6p and Ndt80p at respective promoters (Fig. 4E). We term this arrangement “sequentially convergent promoters.” Thus, *NDJ1* expression is governed by a genetic circuit where temporally overlapping sense and antisense transcription enables dsRNA formation, and where the subsequent dsRNA-triggered vacuolar degradation shapes the observed pulsed pattern.

## Discussion

Cell fate transitions demand the precise expression of stage-specific mRNA cohorts. Equally important, transcripts from earlier stages, which differ in stability and abundance, must be removed to permit proper cell-state progression. The mechanisms that coordinate such broad and synchronous clearance remain elusive. In this study, we identify extensive formation of dsRNA as a key driver of mRNA structural remodeling during early-to-middle meiosis. We show that dsRNA formation is triggered by NATs under the control of stage-specific transcription factors. These dsRNAs assemble into cytoplasmic aggregates and ultimately mediate transcript clearance via the vacuole. This mechanism is both direct and powerful: the same transcription factors that induce mRNAs needed for a specific developmental stage also program the synthesis of NATs that remove transcripts left over from the prior stage. Our findings demonstrate that large-scale dsRNA formation occurs under physiological conditions and establish it as a fundamental pathway that orchestrates selective and timely transcript removal during meiotic stage transitions (Fig. 5).

**Figure 5.**
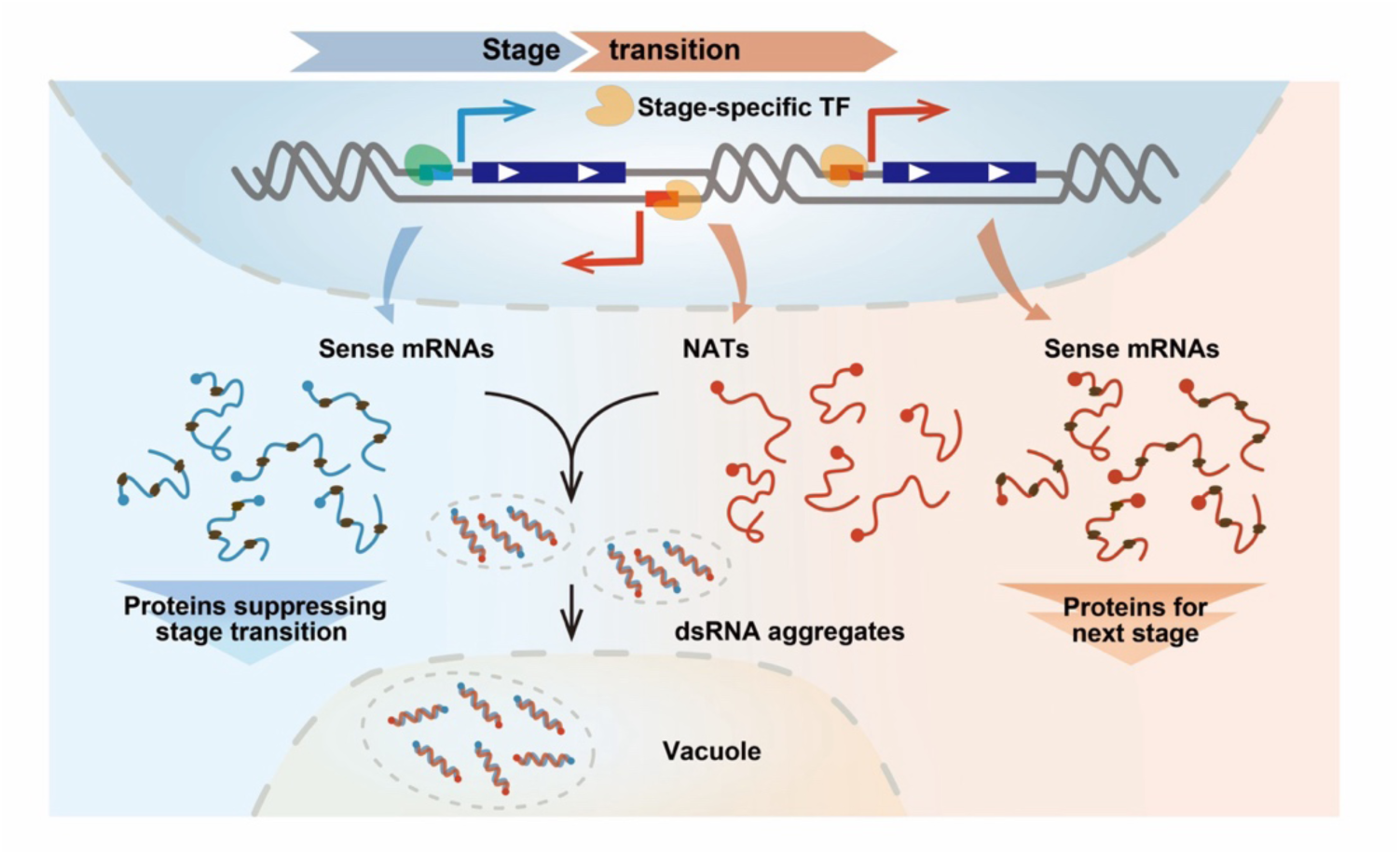
**Programmed dsRNA formation mediates meiotic stage transitions** Model of programmed dsRNA formation driving meiotic stage transitions. Proteins translated from mRNAs of the previous stage act as brakes that prevent premature progression. Stage-specific transcription factors (e.g., Ume6p and Ndt80p) simultaneously induce (*i*) new sense mRNAs required for next stage and (*ii*) pervasive NATs that form dsRNA aggregates with mRNAs that would otherwise inhibit stage transition. These aggregated dsRNAs are rapidly cleared via the vacuolar degradation pathway, thereby enabling progression to the next stage.

We initiated this work by probing RNA structural dynamics across meiosis^21^. Unexpectedly, we traced the most pronounced rearrangements to dsRNA formation. Conceptually, interactions influencing RNA structure can be classified as transient, such as ribosomal elongation, or stable, including persistent RNA–RNA base pairing and certain RNA–protein binding events. Ribosome scanning and elongation have conventionally been viewed as major in-cell disruptors of secondary structure. Our data now show that stable interactions exert large, often greater, and broadly overlooked impacts on the structural changes captured by chemical probing. Comparable chemical probing observations have been made in other biological processes, including human neurogenesis^31^ and zebrafish embryogenesis^32,33^, where pronounced structural changes were primarily attributed to RNA-binding protein or microRNA engagement rather than translational flux. Our study suggests that some structural signatures previously ascribed to ribosomal translocation warrant re-evaluation for potential contributions from alternative stable interactions, especially RNA-RNA interactions.

Remarkably, we identified 245 gene loci associated with antisense transcription during the transition from the pre-meiotic stage (P) to prophase I (T1), which is substantial relative to the roughly 480 total genes that are significantly induced during sporulation^34^. Antisense transcription is thus as pervasive as sense transcription during this transition. These NATs are activated by meiotically essential transcription factors, including Ume6p and Ndt80p. Thus, the same transcription factors simultaneously enable the induction of required new mRNAs and the clearance of pre-existing mRNAs, highlighting their bidirectional regulatory roles. Prior work showed that Ndt80p can up- or down-regulate gene expression through transcriptional isoform switching^28^. Our study uncovers an additional distinct mechanism that achieves comparable regulatory complexity, illustrating another dimension of the multifaceted nature of transcription factor-mediated control.

We observed that dsRNAs form cytoplasmic aggregates. Co-localization analysis with multiple cellular markers suggests these aggregates are delivered to the vacuole for degradation via autophagy. Although a direct mechanistic linkage between dsRNA aggregates and autophagy remains preliminary, several observations strongly support this model. First, large dsRNA aggregates are detected inside the vacuole, and autophagy is the primary route by which bulk cytoplasmic material is delivered there^35^. Additionally, the timing of meiosis-specific antisense transcription aligns with known meiotic functions of autophagy, which is required for entry into the first meiotic division but not the second^36^. The *TIP1* sense/antisense RNA localization analyses confirmed that NATs mediate selective transcript aggregation. Given that NATs encode nonsense transcripts, aberrant translation of NATs could facilitate selective dsRNA aggregation through involvement of nonsense-mediated decay factors such as UPF1^37^. dsRNA sensors may also confer specificity, since selective autophagy of pathogen dsRNA occurs in human cells via direct binding of the cargo receptor SQSTM1 to dsRNA^38^. Defining the precise molecular mechanisms underlying this selective degradation pathway is an important area for future study.

Most broadly, our study emphasizes that transcriptome remodeling requires bidirectional transcription, involving activation of new transcripts and concurrent elimination of obsolete ones, where the latter process has long been underestimated. Antisense transcription appears pervasive during events involving large-scale changes in cell state, including gametogenesis^17,39^, embryogenesis^40,41^, eye development^42^, stress responses^43^, and meiosis. We propose that dsRNA-mediated clearance of sense transcripts may constitute a widespread mechanism for programming cell fate transitions. For example, dsRNAs detected in mouse testes, associated with pachytene-stage genes^17^, parallels our findings in yeast and suggests that dsRNA formation during meiosis is functionally conserved. Given that large-scale dsRNA accumulation can trigger innate immune responses^44^ and that many species possess Dicer-dependent dsRNA cleavage pathways, large-scale dsRNA formation may be limited or preferentially occur in immune-privileged contexts such as the germline and early embryo. Importantly, our findings show that shifts in the relative strengths of convergent promoters enable dsRNA formation. Decoding cis-acting elements within these promoters and their temporal interplay provides a direct strategy to uncover additional programmed, transient dsRNA formation events and to define their regulatory roles.

### Limitations of this study

We note the following limitations that should be considered when interpreting the results. (*i*) Rapid dsRNA-mediated degradation of sense mRNAs, especially at the synchronized pachytene (T0) stage, limits their structural coverage in our dataset. This explains why the most pronounced structural changes are observed during the T0–T1 interval, as accurate RNA structure determination requires high sequencing depth. (*ii*) The digestion efficiency of the USER enzyme used in stranded library preparation is not 100%, meaning that the antisense read ratio contains background signal (∼15%) arising from undigested second-strand cDNA. (*iii*) RNA abundance analyses in this study reflect a combination of transcriptional and post-transcriptional effects and therefore cannot be interpreted as exclusive evidence of post-transcriptional regulation. (*iv*) The MS2/MCP system introduces background signals from MCP-GFP. (*v)* Mapping transcription initiation sites using nanopore full-length cDNA sequencing may reflect both true transcriptional start sites and artifacts arising from *in vivo* RNA degradation or degradation introduced during library preparation.

## Data availability

The raw DMS-MaP and long-read sequencing data are available at the NCBI BioProject under accession number PRJNA1174166. Scripts used for sequencing data processing and analysis are accessible at the GitHub repository: https://github.com/HoNg621/StructuromeYeastMeiosis. Further inquiries regarding resources and reagents should be directed to the lead contact, H.W.

## Acknowledgments

This work was supported by the National Key Research and Development Program of China (2016YFA0500900 to Y.B.), start-up funding from ShanghaiTech University (to Y.B.) and the US National Institutes of Health (R35 GM122532 to K.M.W.). We thank the Molecular and Cell Biology Core Facility (ShanghaiTech University) and the Molecular Imaging Core Facility (ShanghaiTech University) for technical support. We thank Jilong Liu (ShanghaiTech University) and Kun Dou (ShanghaiTech University) for sharing plasmids.

## Author contributions

Y.B. and H.W. conceived of and supervised the study. H.W. and Y.B. designed the experiments. H.W. designed the analyses. H.W. and X.L. performed the experiments with assistance from Y.X. H.W. performed the analyses. H.W. and K.M.W. wrote and revised the manuscript.

## METHODS

### Synchronous meiosis conditions

The SK1-derived yeast strain A14201, frequently used as a model for meiotic studies, was first cultured in YPD medium (1% yeast extract, 2% peptone, 2% glucose) until reaching saturation. Cells were then transferred into YPA medium (1% yeast extract, 2% peptone, 1% potassium acetate, pH 7.0) at an initial OD_600_ of 0.3 and allowed to grow overnight. Cultures were washed twice and shifted into SPO medium (0.5% potassium acetate, pH 7.0, 0.02% raffinose) at an OD_600_ of 1.9 to induce meiosis. Six hours following the switch to SPO, 1 µM β-estradiol was added to induce expression of the meiotic transcription factor Ndt80.

### dsRNA immunofluorescence

Yeast immunofluorescence was performed as described^46^ with modifications. Cells were fixed in 3.7% formaldehyde for 20 minutes at room temperature, washed twice with 1x PBS, and resuspended in 100 μl of 1 M sorbitol/PBS. Cell walls were digested with 0.2 U/μl lyticase (Yeasen, 10403ES81) at 30 °C for 20 minutes for vegetative cells or 30 minutes for sporulating cells. The cells were then adhered to polylysine-coated slides and permeabilized in −20 °C methanol for 6 minutes, followed by acetone for 30 seconds. After air drying, cells were blocked with 5% bovine serum albumin and incubated with anti-dsRNA antibody rJ2 (from mouse; Sigma, MABE1134, 1:60), followed by Cy3-conjugated goat anti-mouse IgG secondary antibody (Jackson ImmunoResearch, 115-165-003, 1:200). Nuclei were counterstained with DAPI.

### Hierarchical clustering

RNA-seq reads were assigned to gene loci in a strand-specific manner using featureCounts^47^ and normalized to TPM values. At each time point, genes with TPM greater than 10 were retained for antisense read ratio analysis, whereas those with TPM below 10 were treated as missing values. Genes with antisense read ratio changes greater than 20% across all time points, with no more than one missing value, were included in the hierarchical clustering. Clustering was performed using Cluster 3.0^48^ with centroid linkage based on centered Pearson correlation, and clustering results were visualized using Java Treeview^49^.

### Northern blotting analysis

Yeast genomic DNA was extracted to serve as a PCR template for probe synthesis. Briefly, 2 ml of cells were lysed in 900 μl genomic DNA lysis buffer (0.2 M LiOAc, 1% SDS), incubated at 70 °C for 20 min, and centrifuged at 15,000 ×g for 5 min. The supernatant was collected, and DNA was precipitated with isopropanol. The resulting DNA was dissolved, treated with RNase A (ThermoFisher Scientific, EN0531) at 37 °C for 30 min, and purified by phenol-chloroform extraction. Northern blot probes were generated by PCR amplification of a ∼500-nt exon fragment of the target gene using primers containing a T7 promoter sequence. PCR products were purified with the EasyPure PCR Purification Kit (TransGen Biotech, EP101-01), and *in vitro* transcription was performed using the DIG Northern Starter Kit (Roche, 12039672910). After transcription at 42 °C for 1 h, RNA probes were diluted to 50 μl with RNase-free water, aliquoted, and stored at −80 °C. Probe labeling efficiency was assessed according to the kit protocol.

For Northern blotting, 2 μg of total RNA was denatured in NorthernMax-Gly loading dye (ThermoFisher Scientific, AM8551) at 50 °C for 1 h and resolved on a 1% agarose gel prepared with 1× PB buffer (10× PB: 8.2 g/L Na₂HPO₄, 5.1 g/L NaH₂PO₄). Electrophoresis was performed at 4 °C (4 V/cm) with periodic buffer replacement. RNA was transferred to a nylon membrane by capillary blotting in 20× SSC for >6 h and crosslinked by UV irradiation (200 mJ, four times). Membranes were prehybridized in ULTRAhyb-Oligo buffer (ThermoFisher Scientific, AM8663) at 68 °C for 30 min, followed by overnight hybridization with heat-denatured RNA probes. Post-hybridization washes and detection were performed according to the DIG Northern Starter Kit protocol.

### RNA and protein tagging for localization analysis

The pRS shuttle vector, which contains an autonomously replicating sequence and a centromeric sequence, was used to exploit endogenous replication and chromosome segregation machinery in yeast.

For RNA tagging, a cassette of 24×MS2 stem–loop repeats was inserted upstream of the *ADH1* terminator in a pRS vector, carrying a G418 resistance marker. The ORF of the target gene, together with its native 1 kb promoter, was cloned immediately upstream of the 24×MS2 repeats to generate MS2-tagged transcripts. The MCP-GFP fusion ORF was cloned under the control of the *NFS1* promoter into a separate pRS vector carrying a nourseothricin resistance marker. MS2-tagged and MCP-GFP plasmids were co-introduced into yeast cells by electroporation, and transformants were selected on medium containing G418 and nourseothricin. Verified colonies were induced to undergo synchronous meiosis, and samples were collected at designated time points. Cells were fixed in 3.7% formaldehyde for 20 min at room temperature for dsRNA immunofluorescence and co-localization analysis.

For protein tagging, the ORF of each gene together with its native 1 kb promoter was cloned into a pRS vector carrying a G418 resistance marker. GFP was fused to the C terminus of Edc3, Pab1, and Vma2, and to the N terminus of Atg8. The tagged plasmids were transformed into yeast cells, and transformants were selected for induction of synchronous meiosis. Cells were fixed in 3.7% formaldehyde for 20 min for dsRNA immunofluorescence and co-localization analysis.

### Motif enrichment analysis

One-kilobase downstream regions for genes with meiosis-specific antisense transcription were retrieved from annotation files using a custom script. Equivalent regions from other genes were retrieved as the background set. Motif enrichment analysis was performed using STREME^50^ with parameters --dna --minw 6 --maxw 10 --nmotifs 8. Transcription factors corresponding to enriched motifs were identified with Tomtom^51^, using the enriched motifs as input. Binding motifs of identified transcription factors were downloaded from the JASPAR database^45^ and used as input for FIMO^52^ to map their precise locations in antisense promoters. High-scoring promoters were selected for subsequent experimental validation.

### Long-read sequencing and ChIP-seq data processing

For long-read sequencing, libraries were prepared using the SQK-PCS109 and SQK-PBK004 kits (Oxford Nanopore Technologies) according to the manufacturer’s protocol. Briefly, 500 ng of total RNA was used for reverse transcription and strand switching with Oligo(dT) VN primers and strand-switching primer. The reverse-transcribed cDNA was amplified and purified using AMPure XP beads (Beckman Coulter, A63881). Purified cDNA was ligated with adapters. Libraries were loaded onto an R9.4 flow cell and sequenced on a PromethION platform (Oxford Nanopore Technologies).

Sequences were base-called from raw fast5 files using Guppy (v5.0.16). Reads were aligned to the reference genome using Minimap2 (v2.1.1-r341)^53^ with parameters -ax splice -uf -k14. The resulting BAM files were sorted and separated by strands with SAMtools^54^. The 5′ end locations were extracted and merged into BED files for mapping transcription initiation sites using a custom script. The stranded BAM and BED files were visualized directly in IGV^55^.

The Ume6 and Ndt80 ChIP-seq raw reads were downloaded and aligned to the reference genome by BWA^56^. The resulting BAM files were sorted and used for peak calling with MACS2^57^. Identified peaks and corresponding BAM files were visualized directly in IGV^55^.

### Reverse transcription quantitative PCR

Total RNA was isolated from meiotic cells using a modified hot phenol method^58^, followed by column purification with the TransZol Up Plus RNA Kit (TransGen, ER501-01). The purified RNA was treated with RNase-free DNase I (Promega, M6101) to remove genomic DNA. For cDNA synthesis, 100 ng of total RNA was reverse-transcribed using the HiScript III All-in-One RT SuperMix Perfect for qPCR kit (Vazyme, R333-01). Quantitative RT-PCR was performed for the indicated genes using TB Green Premix Ex Taq™ II (Takara, RR820A) on a QuantStudio™ 7 Flex Real-Time PCR System. Relative expression levels were normalized to *ACT1* and calculated using the –ΔΔCt method.

### Statistics and reproducibility

Unless specified, statistical tests were performed using the SciPy v.1.13 package in Python. Correlation analyses between DMS reactivities used Pearson’s correlation. The number of data points for each analysis is indicated in the figure legends. The long-read sequencing experiment was performed once. Other experiments were repeated at least three times with consistent results.

## SUPPLEMENTAL FIGURE TITLES AND LEGENDS

**Figure S1.**
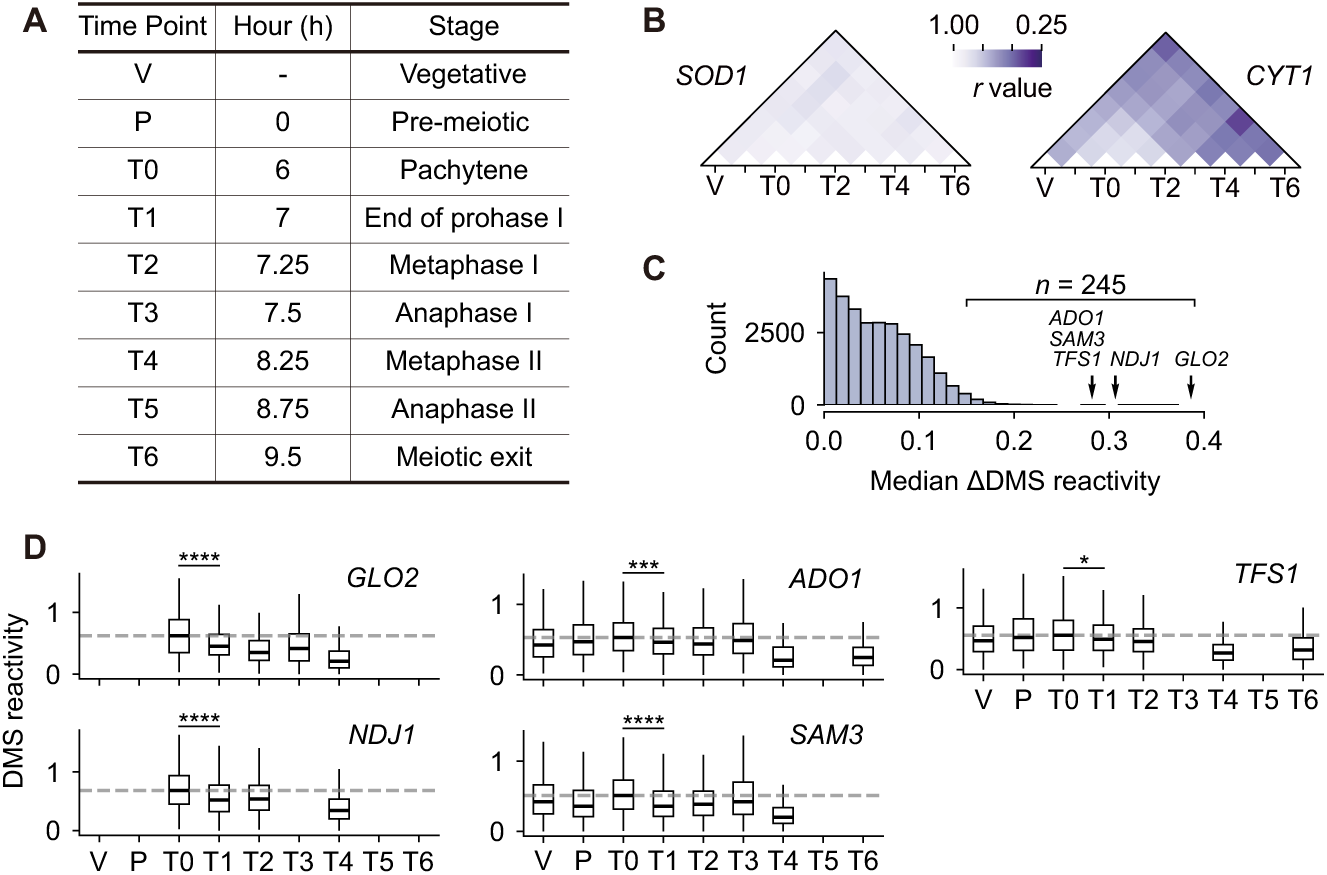
Time course of synchronized yeast meiosis and analyses on DMS reactivity changes across stages, related to. **Figure 1** (A) Meiotic time points and corresponding stages in the synchronized Ndt80-inducible system. Hours represent time in SPO medium after Ndt80 induction at T0 (6 h). Staging is as defined in our previous study^21^. (B) Examples of conserved (*SOD1*) and strongly remodeled (*CYT1*) mRNA structures during meiosis. (C) Distribution of median ΔDMS reactivities for 28,601 pairwise comparisons. (D) Distribution of DMS reactivity at each stage for *GLO2*, *NDJ1*, *ADO1*, *SAM3*, and *TFS1*. For all five genes, DMS reactivity drops significantly between T0 and T1 and reaches its maximal decrease by T4 (median change shown). The dashed line indicates the median DMS reactivity at T0.

**Figure S2.**
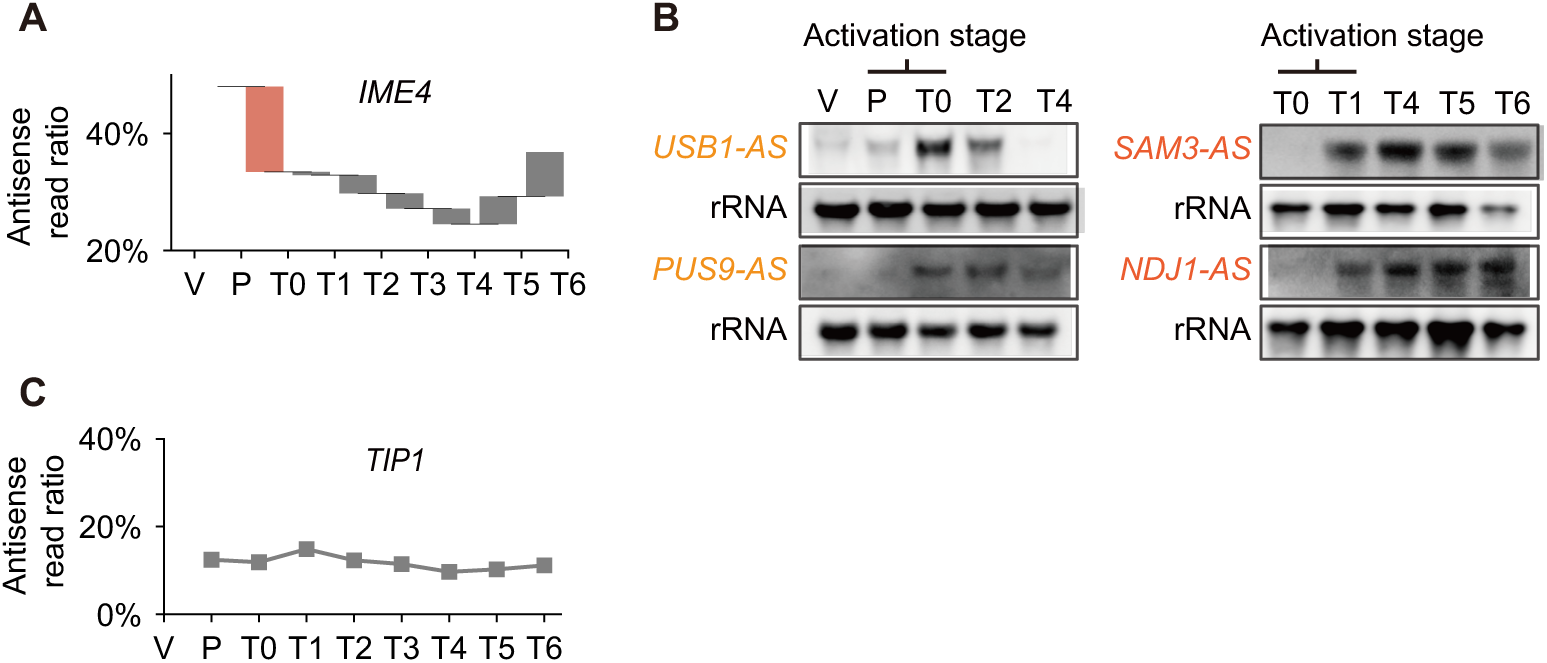
Analyses on antisense read ratios and validation of antisense RNAs by northern blot, related to. **Figure 2** (A) Antisense read ratio of *IME4* throughout meiosis. The orange segment highlights the interval with the largest change. (B) Northern blot analysis of antisense RNAs: *USB1-AS*, *PUS9-AS*, *SAM3-AS*, and *NDJ1-AS*. The activation stage is defined as the interval with the largest increase in antisense read ratio. (C) Antisense read ratio of *TIP1* throughout meiosis. The ratio remains stable at a background level of 15%–20%.

**Figure S3.**
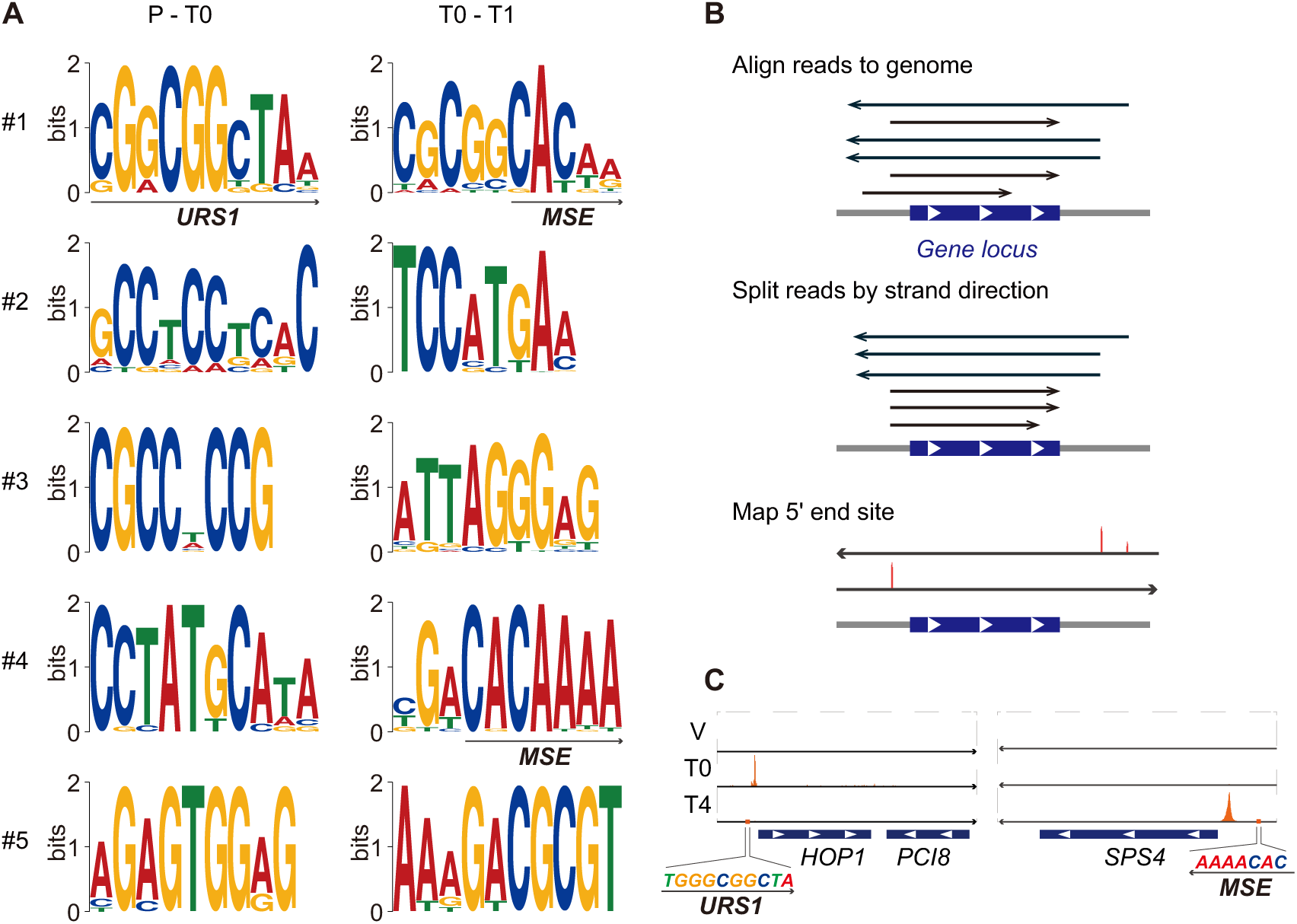
Motif enrichment analysis results and overview of 5′ end mapping by long-read sequencing, related to. **Figure 3** (A) Top five motifs enriched in the motif enrichment analyses for antisense promoters activated in the P-T0 and T0-T1 intervals. Core sequences of the URS1 (Ume6-binding) and MSE (Ndt80-binding) elements are highlighted within the respective motifs. (B) Overview of long-read sequencing and identification of transcription initiation sites based on 5′ end mapping. Transcription initiation sites are indicated by the orange peaks. (C) Transcription initiation sites (orange peak) for *HOP1* and *SPS4* sense mRNAs at the V, T0, and T4 stages.

**Figure S4.**
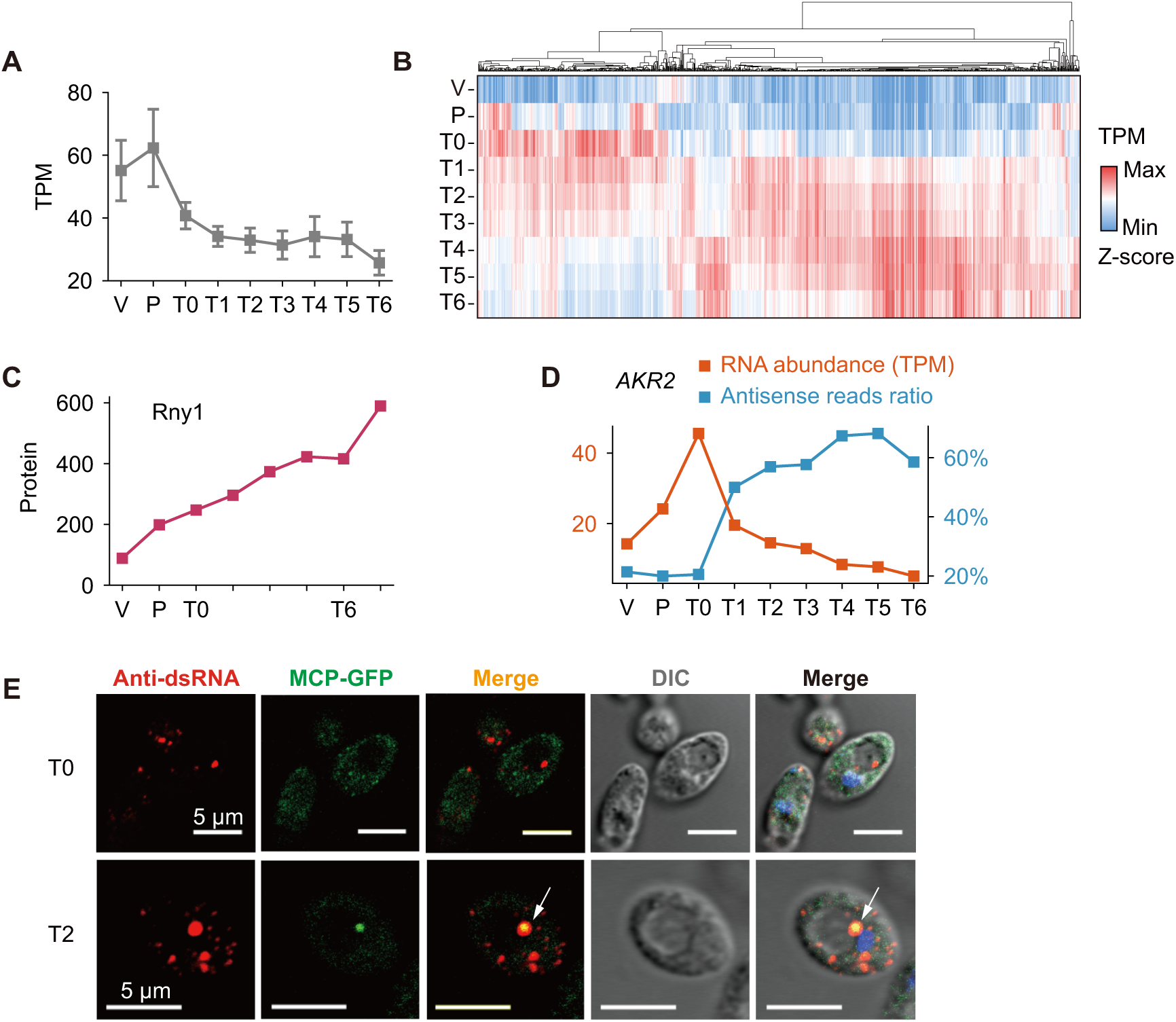
Sense mRNA profiles, Rny1p abundance across meiosis and *AKR2* mRNA localization analysis, related to. **Figure 4** (A) Average RNA abundance of 313 genes exhibiting meiosis-specific antisense transcription. (B) Hierarchical clustering of RNA abundance for 1,044 transcripts that were upregulated more than eightfold during meiosis and did not exhibit meiosis-specific antisense transcription. (C) Protein abundance of Rny1 throughout meiosis detected by mass spectrometry^28^. (D) Relationship between *AKR2* RNA abundance and antisense read ratio. (E) Co-localization analysis of *NDJ1-MS2* with dsRNA aggregates before (T0) and after (T2) Ndt80 induction. White arrows indicate co-localization between dsRNA signals and MCP-GFP.

## Supplemental information

Table S1. Pearson correlation analysis of DMS reactivities and median ΔDMS reactivity for 28,601 pairwise comparisons across 2,058 mRNAs, related to Fig. 1A and 1B.

Table S2. Antisense read ratios for 5,520 gene loci across all time points. Genes with changes greater than 20% and no more than one missing value were used for hierarchical clustering, related to Fig. 2A.

Table S3. Genes with meiosis-specific NATs (*n* = 313) classified by meiotic stage of maximum increase in antisense read ratio, related to Fig. 2B and 2C.

Table S4. Motif enrichment analysis results for antisense promoters of genes classified into P-T0 and T0-T1 intervals, related to Fig. 3A and S3A.

Table S5. Raw results of RT-qPCR for MSE deletion experiments, related to Fig. 3E.

Table S6. Hierarchical clustering results of sense mRNA abundance profiles for genes associated with meiosis-specific NATs, related to Fig. 4A.

Table S7. Peak calling results for Ume6p and Ndt80p binding sites, related to Fig. 4E. The ChIP-seq data used in this analysis are from previously published datasets^29,30^.

